# LI-Seq: A Cost-Effective, Low Input DNA method for Whole Genome Library Preparation

**DOI:** 10.1101/2021.07.06.451326

**Authors:** Teia M. Schweizer, Matthew G. DeSaix, Kristen C. Ruegg

## Abstract

1. Samples from species of high conservation concern are often low in total genomic DNA. Whole Genome Sequencing (WGS) can provide many insights that can be used to aid in species conservation, but current methods for working with low quality and low input samples can be cost prohibitive for population level genomic analyses. Thus, there is an urgent need for a cost-effective method of preparing WGS libraries from low input DNA samples.
2. To bridge the gap between sampling techniques commonly used in conservation genetics that yield low quality and low input DNA and the powerful tool of WGS, we developed LI-Seq, a more efficient method that successfully produces libraries from low quality DNA with as low input as 0.48 ng of DNA, with an average final library size of 300-500 base pairs.
3. Sequencing results suggest no difference in sequencing quality or coverage between low quality, low input and high quality, high input starting material using our protocol. We conclude that our new method will facilitate high-throughput WGS on low quality, low input samples, thus expanding the power of genomic tools beyond traditional high quality samples.

## Introduction

The field of genomics began developing in the 1990s with whole genome shotgun sequencing of bacteria (Weissenbach 2016) and has advanced rapidly with the improvement of High-Throughput Sequencing (HTS) techniques (Goodwin et al 2016). The ability to sequence whole genomes with relative ease has opened new research avenues and made it possible to estimate fundamental population genetic parameters with increasing precision in both model and non-model organisms (Allendorf, Hohenloe, and Luikart 2010; Ouborg et al 2010). Of particular relevance to ecologists and conservation biologists, HTS has made it possible to investigate previously challenging topics such as the genetic basis of local adaptation, patterns of inbreeding across the genome, and how species adapt to changing climate conditions (Kohn et al 2006; Ruegg et al 2018; Allendorf, Hohenloe, and Luikart 2010; Ouborg et al 2010). As a result, genomic tools are revolutionizing the fields of ecology, evolution, and conservation biology.

Despite the proliferation of HTS methods for model organisms (Ekblom and Galindo 2011), there remain a number of technical and financial limitations to the widespread use of genomic approaches in situations where the amount of input DNA maybe limited and costs are a concern. While the cost of sequencing has dramatically decreased in the last two decades (Goodwin et al 2016), it is often still prohibitively high for use in population-level studies where hundreds or thousands of individuals must be sequenced (Fuentes-Pardo and Ruzzante 2017). Methods modified from commercially available whole genome sequencing (WGS) library preparation kits offer low coverage options at a fraction of the cost per individual, making them suitable for population genetics studies (Therkildsen and Palumbi 2017; Kryazhimskiy et al 2014; Baym et al 2015). However, these methods still typically require high quality and high input DNA and are not optimized to efficiently amplify smaller target library sizes. Such high quality and quantity DNA can be difficult to attain when working with threatened, endangered, or cryptic species, where ethical and logistical challenges are often prohibitive (Kohn et al 2006; Ouborg et al 2010). However, samples that yield low quality and quantity DNA have previously found limited use in whole genome studies unless potentially cost-prohibitive library preparation kits or methods are employed (Taylor et al 2020). Given the immense potential benefits of analyzing whole genomes for effective wildlife conservation and management efforts (Funk et al 2012; Ryder 2005; Russello et al 2015), there is an urgent need for a cost-effective method of preparing WGS libraries from low quality and quantity DNA samples.

Low quality and quantity samples are often a hallmark of noninvasive or minimally invasive sampling techniques. Noninvasive genetic sampling methods first gained recognition in 1992 when DNA was successfully extracted from passively-collected hair for a genetic study of an endangered bear species (Taberlet and Bouvet 1992). Since then, noninvasive genetic sampling has been successfully used in genetic studies across myriad taxonomic groups (Stenglein et al 2010; Valiere et al 2003; Roques et al 2014; Regnaut et al 2006). Noninvasive sampling encompasses samples such as saliva, hair, feces, or feathers, collected without capturing, handling, or otherwise disturbing the study organism (Waits et al 2005). Minimally invasive sampling entails capturing or handling a study organism with minimal invasion or tissue collection (e.g. feather pulls and buccal swabs) (Carroll et al 2017). In recent years, noninvasive and minimally invasive sampling methods have gained popularity, especially for use in monitoring threatened and endangered species (Lukacs and Burnham 2005; Fuentes-Pardo and Ruzzante 2017). However, noninvasively or minimally invasively collected samples typically yield lower concentrations of DNA which can limit their use in whole genome studies unless expensive library preparation kits or library preparation methods are employed (Taylor et al 2020). Given the immense potential benefits of analyzing whole genomes for effective wildlife conservation and management efforts (Funk et al 2012; Ryder 2005; Russello et al 2015), there is an urgent need for a cost-effective method of preparing WGS libraries from noninvasively or minimally invasively collected samples.

Although low cost methods for WGS exist (Therkildsen and Palumbi 2017; Kryazhimskiy et al 2014; Baym et al 2015), they still require a prohibitively large amount of high quality input DNA (2.5 ng) for many conservation applications. More specifically, further analysis of these methods reveals that much of the DNA is wasted during the library preparation step due to the fact that the average fragment size produced from these methods is 1kb, but the average fragment size needed for many common sequencing platforms, such as Illumina, is 300-500 base pairs. Thus, DNA above 500 base pairs is often removed prior to sequencing. To bridge the gap between sampling techniques commonly used in conservation genetics that yield low quality DNA and the powerful tool of WGS, we developed a more efficient method that successfully produces libraries from low quality DNA with as low input as 0.48 ng of DNA, with an average final library size of 300-500 base pairs.

We demonstrate the utility of our method for producing high quality sequencing data at a fraction of the cost of traditional library preparation methods using DNA extracted from a single flight feather calamus, or quill, of a small (8-9 grams) passerine bird, the American Redstart (*Setophaga ruticilla*). We compare the sequence data from our low input DNA library (from feather) to those generated from a high input library (from blood) and demonstrate that our method produces comparable sequence quality for both low and high input DNA sources. These results have important implications for conservation genomics research seeking to maximize efficient sequencing from low input DNA samples.

## Materials and Methods

### Library Preparation for Sequencing

We identified three key parameters in other methods (i.e. Therkildsen and Palumbi 2017) that we could optimize in order to target the ideal fragment distribution and avoid loss of critical DNA when working with low input DNA samples. The three key parameters modified herein were: (1) the ratio of tagmentation transposome (which cleaves DNA and adds an adapter for indices) to input DNA quantity, (2) the duration of tagmentation incubation, and (3) the duration of the indexing PCR elongation time.

In order to compare sequencing quality from high and low input and quality DNA libraries and assess the efficiency of our method, we extracted DNA from 50 high DNA quantity and quality bird blood samples and 50 low DNA quantity and quality feather calamus tips of the American Redstart. For blood, DNA was extracted from between 50-100 μL of whole blood stored in Queen’s Lysis Buffer (∼80 μL of whole blood plus 300 μL of buffer), using Qiagen DNEasy Blood and Tissue Kit and eluted into 100 μL of provided AE buffer. For extractions from feathers, like other low-quality samples, maximizing DNA yield is critical. Therefore, we followed the Qiagen protocol but with the following modifications. To each sample, we added 10 μL of 1 M Dithiothreitol (DTT) to the initial lysis step to aid in breaking down disulfide bonds found in the keratin of feathers. Flowthrough after the first filtration step when lysate was transferred to the spin column was pipetted back onto the filter for a second centrifugation. Prior to the final elution step, AE buffer was placed in an incubator at 56 °C. During the final elution step, AE buffer was left to incubate on the filter for five minutes instead of two. We eluted feather extractions into 400 μL (two rounds of 200 μL elutions through the spin column as recommended by Qiagen protocol for maximum yield). Prior to proceeding with library prep, we concentrated feather DNA extractions using a 1:1 ratio of Serapure beads (Faircloth and Glenn 2014) from 400 μL to 15 μL and eluted into 10mM Tris-Hcl (Figure 1, step 1a).

**Figure 1.**
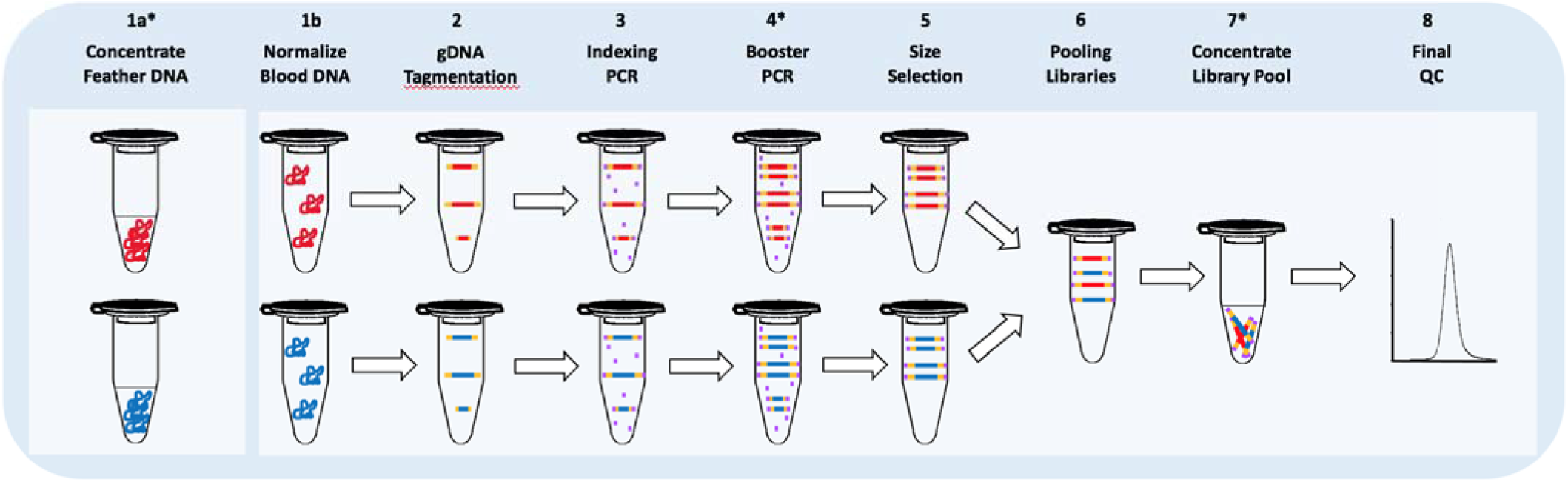
Lab workflow diagram for LI-Seq method. Steps 1 through 8 follow full protocol (see Supplementals). Steps with * after number denote steps where modifications to typical WGS library preparation methods were implemented. Low input samples start with step 1a and high input samples start with step 1b.

To ensure that final library concentrations will be similar across samples, we quantified each DNA extraction using a Qubit dsDNA High Sensitivity Assay Kit (Invitrogen) and normalized each sample to a concentration of 0.48 ng/μL-4.5 ng/μL, with a target of 2.5 ng/μL (Figure 1, step 1a and 1b). To fragment the DNA and “tag” it with Nextera adapters, we added 2.50 μL of TD Buffer and 0.5 μL of TDE1 Enzyme (Illumina) to 1 μL of normalized DNA and incubated the samples in a thermocycler at 55 °C for 20 minutes (Figure 1, step 2).

To amplify the tagmented DNA and add Nextera indexing adapters for sequencing, we pipetted 1 μL of each index primer into the appropriate well of tagmented DNA until all samples had a unique pair of dual indexes. We then added 6.0 μL of Kapa Hifi Hotstart Mix (KMM; Kapa Biosystems) before running in a thermocycler as follows: held at 72°; for 3 minutes, held at 98° for 2 minutes and 45 seconds, cycled 8 times through 98° for 15 seconds, then 62° for 30 seconds, then 72° for 30 seconds, held at 72° for 1 minute, and then held at 4° until removed from thermocycler (Figure 1, step 3). As per the Therkildsen and Palmubi (2017) method, this indexing PCR had more cycles than the original Illumina protocol and was broken into two stages (Indexing PCR and “Reconditioning PCR, ” which was renamed Booster PCR in this method). Additional cycles were added in the Therkildsen and Palumbi (2017) method because the tagmented DNA was not purified prior to Indexing PCR, making the PCR reaction less efficient. To further amplify, or “boost,” copies of indexed DNA without using additional Nextera indices, we added 7.6 μL of KMM, 4.4 μL of ultrapure water, and 1.6 μL each of a custom 10uM primer pair (P1=AATGATACGGCGACCACCGA; P2=CAAGCAGAAGACGGCATACGA) to each library. We ran the samples in a thermocycler as follows: held 95 for 5 minutes, cycle 4 times through 98° for 20 seconds, then 62° for 20 seconds, then 72° for 2 minutes, hold at 72° for 2 minutes, and then held at 4° until removed from the thermocycler (Figure 1, step 4).

To purify the PCR product and remove undesirable fragments, we followed standard Ampure bead protocol (Beckman Coulter) using a 0.7:1 bead to DNA ratio which will remove below approximately 320 bp and eluted into 30 μL of 10mM Tris-Hcl (Figure 1, step 5). In order to avoid overrepresentation of one individual during whole genome resequencing, we then quantified using a Qubit dsDNA High Sensitivity Assay Kit (Invitrogen) and pooled an equal number of copies of each sample into a 1.5 mL tube (Figure 1, step 6). Finally, in order to increase the final concentration of the pooled libraries and increase sequencing efficiency, we then followed the standard Ampure double-sided size selection protocol, using a 0.63:1 bead to DNA ratio to remove large fragments and a 0.73:1 bead to DNA ratio to remove small fragments, and eluted into 30 μL of 10mM Tris-HCl (Figure 1, step 7). After the pooled library has been concentrated and double size selected (either with or without the optional reconditioning PCR), we perform final quality control (QC) with Qubit quantification and Tapestation 2200 fragment distribution analysis (Agilent) (Figure 1, step 8).

To address issues of overamplification, also called ‘PCR bubble,’ we encountered while using Therkildsen and Palumbi’s original method (2017) with our low input DNA, we added an optional ‘reconditioning PCR’ step which provides additional reagents, especially primers, so that the PCR product does not anneal to itself (Thompson et al 2002). To recondition a final pooled library with overamplification, we added 12μL of pooled, size selected library and added 7.6 μL of KMM, 1.6 μL of 10uM P1 (AATGATACGGCGACCACCGA), 1.6 μL of 10uM P2 (CAAGCAGAAGACGGCATACGA), and 4.4 μL of ultrapure water. We then ran it in a thermocycler as follows: held 95° for 5 minutes, cycled once through 98° for 20 seconds, then 62° for 20 seconds, then 72° for 2 minutes, held at 72° for 2 minutes, and then held at 4° until removed from the thermocycler. Next, we used Ampure beads to clean up and performed an additional double size selection and bead cleanup before quantifying and running the library through a Tapestation 2200 fragment distribution analyzer (Agilent) for final quality control. This reconditioning PCR is optional and may not be required for all library preparations.

Using the above method, we prepared two WGS libraries from American Redstart (*Setophaga ruticilla*) samples for low (2x) coverage sequencing. One library was prepared with 50 unique blood samples of normalized DNA concentrations between 1.18 - 4.78 ng/uL. The other library was prepared with 50 unique feather samples, extracted from a single feather calamus, with starting DNA concentrations of 0.48 - 5.7 ng/μL. Both libraries had one cycle of reconditioning PCR performed on the final library. Libraries were each sequenced on one full 2 × 150 bp PE (paired end) HiSeq 4000 lane (Illumina).

### Bioinformatic Analysis and Quality Checking

We trimmed the sequence data to remove potential PCR artifacts using the program FastUniq version 0.11.9 (Xu et al 2012). PCR duplicates need to be removed in order to ensure high-quality sequence data in downstream processes such as creating scaffolds in whole-genome sequencing. We mapped reads to an assembled genome of the yellow warbler (Setophaga petechia; Bay et al 2018), using the Burrows-Wheeler Aligner software version 0.7.17 (Li and Durbin 2010). The resulting SAM files were sorted, converted to BAM files, and then indexed using samtools version 1.9 (Li et al 2009). Depth of sequencing coverage at every read position was calculated using the depth function in samtools (Li et al 2009). The quality of the BAM files for the two different libraries was assessed by comparing the average read depth by individual as well as the average read depth by scaffold. We quantified the GC content of 100 base pair windows in the BAM files from the two libraries using CollectGcBiasMetrics function in Picard version 2.23.1 (Broad Institute 2019). We determined the proportion of reads that passed quality filters for the two libraries using CollectWgsMetrics in Picard (Broad Institute 2019). Two-tailed t-tests were used to compare the quality diagnostics for the different libraries and were implemented in R version 3.6.2 (R Core Team 2019).

## Results

### Library Preparation for Sequencing

A comparison between libraries prepared using the method of Therkildsen and Palumbi (2017) and our modified method revealed that doubling the ratio of tagmentation enzyme to DNA, increasing the tagmentation time to 20 minutes, and decreasing the indexing PCR elongation time to 30 seconds resulted in maximization of fragments in the target distribution (Table 1; Figure 2).

**Table 1.**
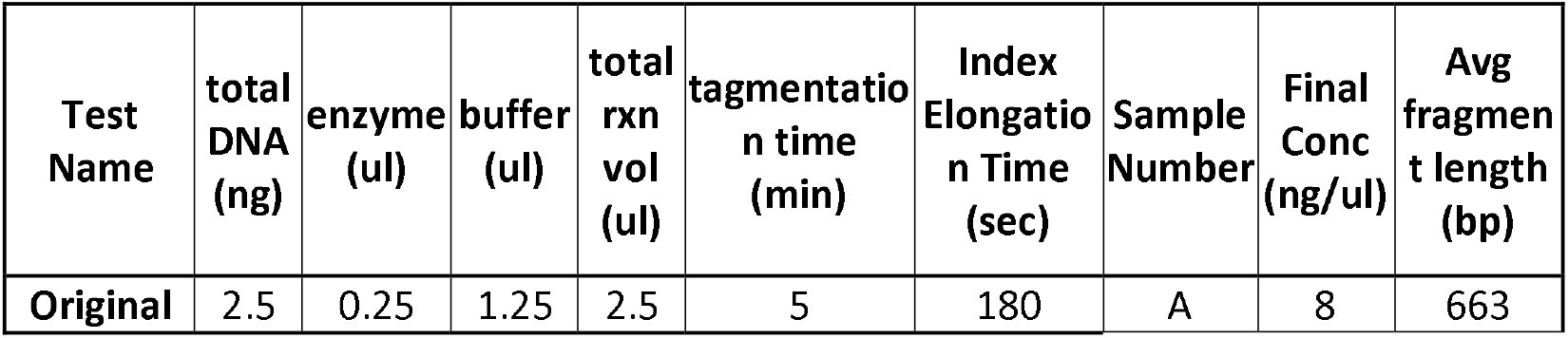
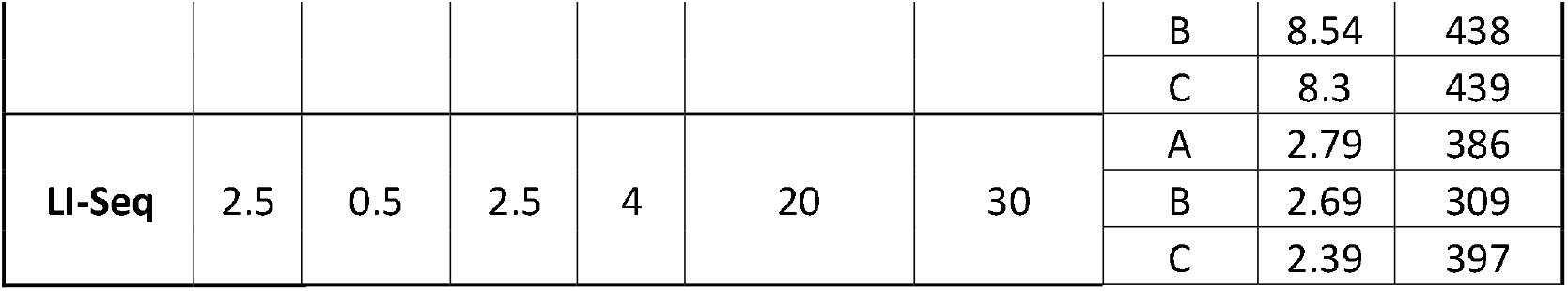
Matrix of the original (Therkildsen) protocol parameters and final LI-Seq parameters, tested with 3 high input samples (blood).

**Figure 2.**
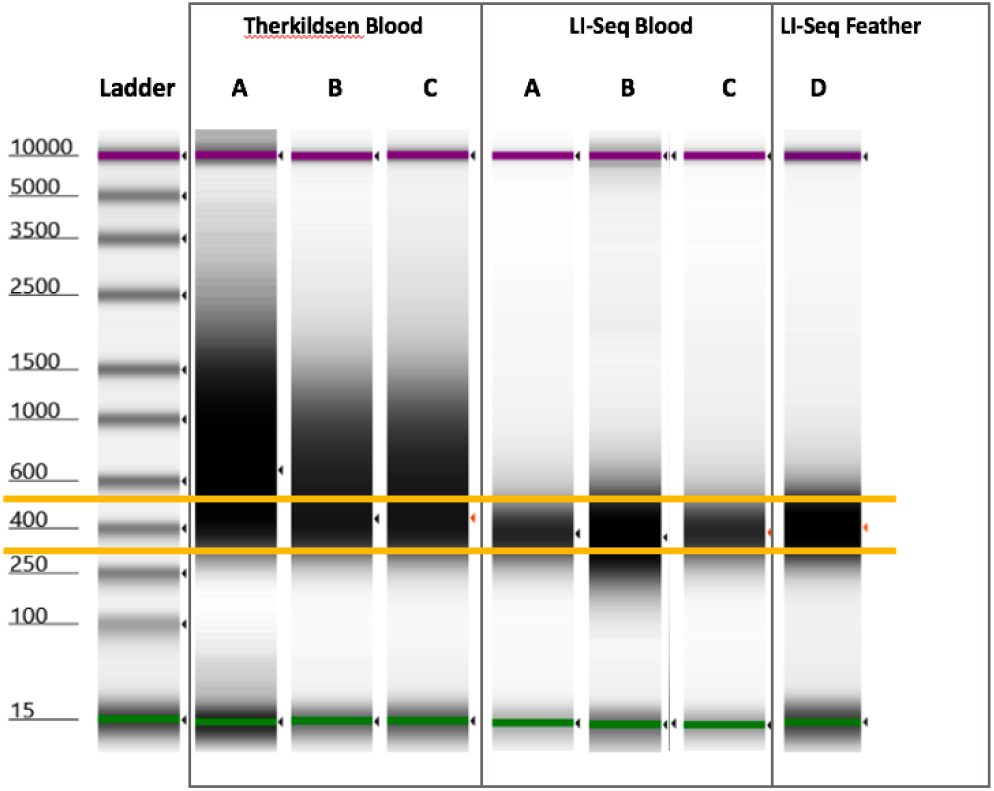
Tapestation 2200 gel of Therkildsen protocol conditions and final LI-Seq conditions from Table 1. Yellow bands show preferred fragment range (320-500bp) for HiSeq 4000 as recommended by Novogene. Individuals A, B and C were duplicated between the Therkildsen Blood and LI-Seq Blood, with the conditions described in Table 1. Individual D was feather DNA prepared using only the LI-Seq method.

Using these modified methods, we successfully prepared two libraries from American Redstart DNA. The library from 50 high input samples (blood) had a final concentration of 12.1 ng/μL, a molarity of 39.3 nM, and an average library size of 466 bp (Figure 3d). The library from 50 low input samples (feathers) had a final concentration of 2.56 ng/μL, a molarity of 8.56 nM, and an average library size of 453 bp (Figure 3f).

**Figure 3.**
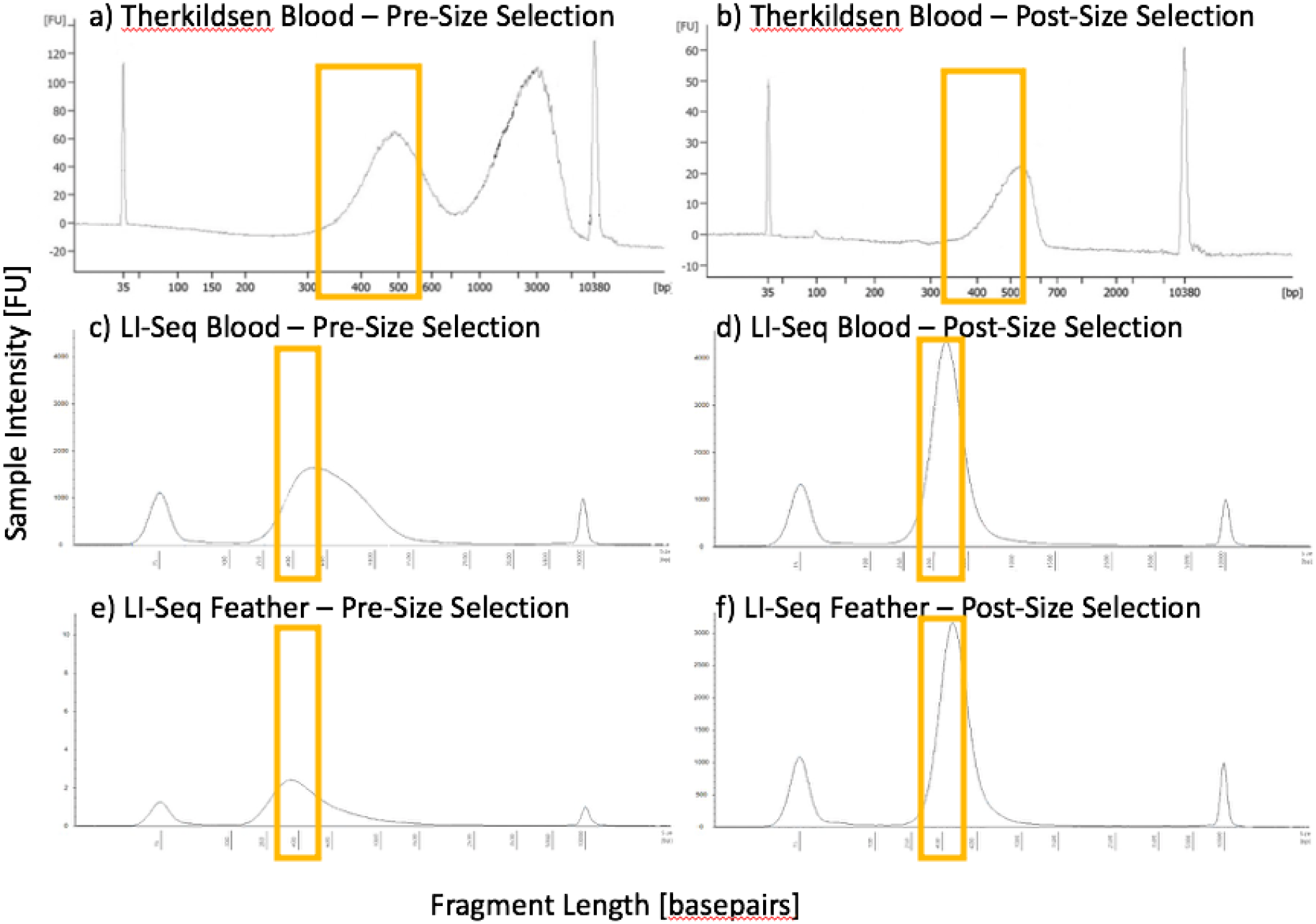
Final library fragment analysis of a library prepared using the original Therkildsen and Palumbi protocol (2017), before (a) and after (b) size selection, run on a Bioanalyzer (Agilent). The second peak of (a) indicated overamplification. The final libraries of the high input (blood) and low input (feather) samples were run on a Tapestation 2200 (Agilent) prior to size selection (high input (c) and low input (e)) and after size selection (high input (d), low input (f)). The x-axis is the size of the DNA fragment in basepairs, and the y-axis is the sample intensity which is correlated with sample concentration. Note the different x- and y-axis scales. Yellow boxes indicate the desired fragment size of ∼320 – 500 bps.

### Bioinformatic Analysis and Quality Checking

The depths of coverage of the sequence data from the two libraries were not significantly different (t = 1.06, df = 98, p-value = 0.29), but the high input library had a slightly higher depth (mean 1.79, standard deviation 0.52; Figure 3a) than the low input library (mean 1.70, standard deviation 0.38; Figure 3a). The high input library had individuals that had a slightly higher proportion of the genome with sequence data (mean 0.66, standard deviation 0.06; Figure 3b) than the low input library (mean 0.64, standard deviation 0.08; Figure 3b), but this difference was not statistically significant (t = 1.85, df = 98, p-value = 0.07). The difference in GC distribution between the two libraries was non-significant (42.9%; t = 1.94e-14, df = 100, p-value = 1), with the mean GC content of the high input library being 42.7% and 42.9% for the low input library (Figure 4C). The sequence data from these two libraries also had very similar patterns of coverage across the scaffolds in the genome (Figure 5). The similarity in coverage across scaffolds shows that certain genomic regions are not being over- or under-amplified in either of the libraries due to a difference in quality and quantity of input DNA. All BAM quality metrics for the two libraries produced comparable results (Table 2).

**Figure 4.**
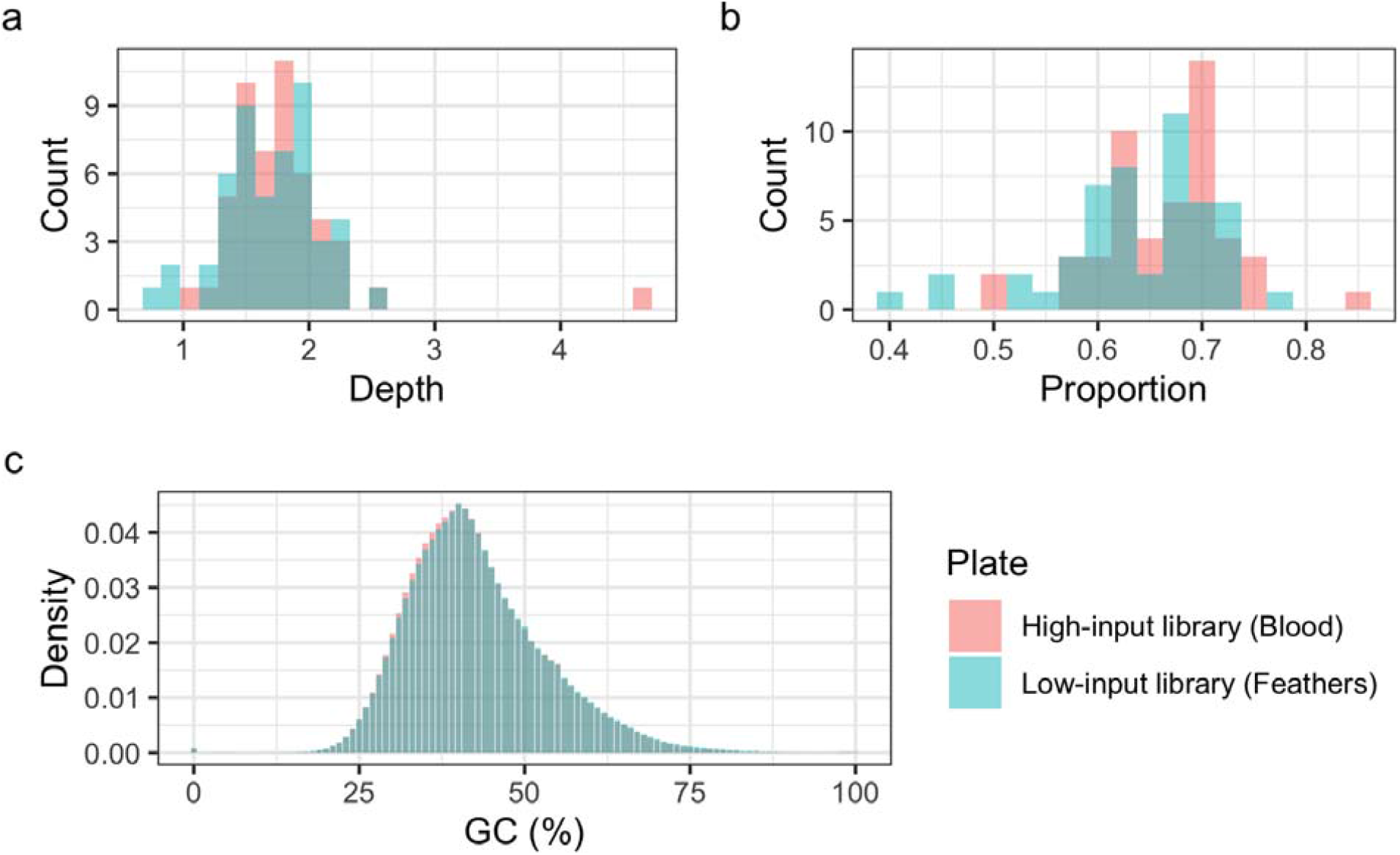
Both histograms of (a) average depth across the genome by individual and (b) proportion of genome with sequence data by individual overlapped considerably between the two libraries and neither measures were significantly different (depth: t = 1.06, df = 98, p-value = 0.29; proportion: t = 1.85, df = 98, p-value = 0.07). GC content was determined for 100 base pair region windows in each individual and then averaged across individuals for each of the two plates. (c) The bins of GC content (ranging from 0 to 100%) were nearly identical for the two libraries (t = 1.94e-14, df = 100, p-value = 1). Data from the high input library from blood samples are shown in red, the low input feather library in light blue, and the overlapping data in darker blue.

**Figure 5.**
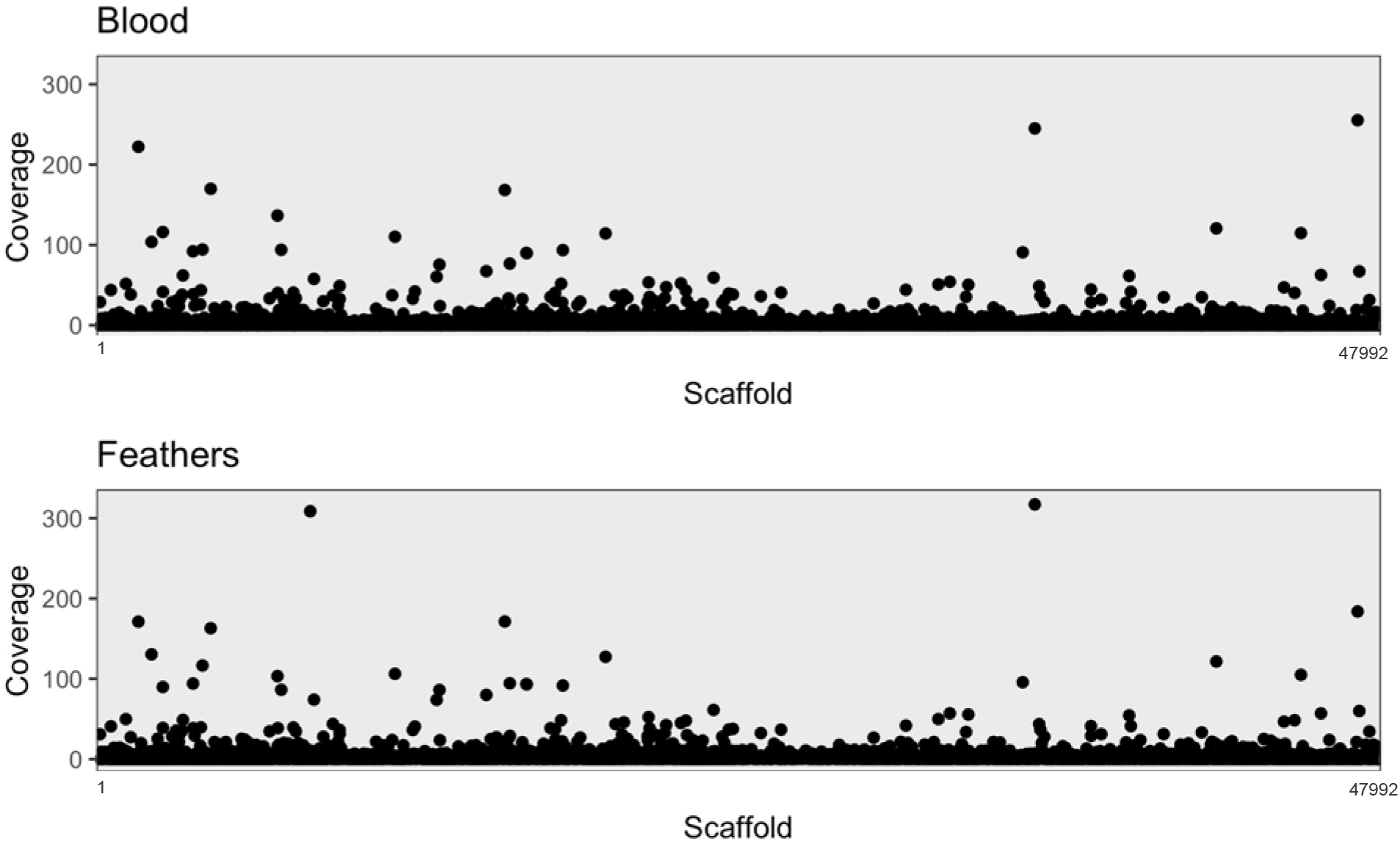
Mean coverage by scaffold averaged across all individuals for the two libraries reveal similar patterns in coverage across the genome between the two libraries.

**Table 2.** BAM quality statistics: the average proportion of reads with adaptors (Adaptors), mapping quality less than 20 (MapQ), marked as duplicates (Duplicate), without a mapped mate pair (Unpaired), quality score less than 20 (BaseQ), and at least 1x (1x) and 5x (5x) coverage after removing low quality reads.

| Library | Adaptors | MapQ | Duplicate | Unpaired | BaseQ | 1x | 5x |
| --- | --- | --- | --- | --- | --- | --- | --- |
| High input<br>(Blood) | 0 | 0.110 | 0 | 0.001 | 0.017 | 0.685 | 0.049 |
| Low input<br>(Feathers) | 0 | 0.117 | 0 | 0.001 | 0.015 | 0.657 | 0.039 |

## Discussion

In conservation genetics, researchers must often work from low quality and low input DNA samples, especially in the case of noninvasive or minimally invasive sampling. These types of samples can be a challenge to use with many genomic techniques, such as WGS, which typically require high input and quality input DNA. Yet, collecting high input samples (like blood and tissue) can present ethical, logistical, and financial roadblocks.

The potential knowledge gained from utilizing WGS and similar genomic tools can provide insights not yet achievable from other methods. WGS can be a powerful tool for management in conservation efforts but, until now, has been challenging and or cost-prohibitive when working with low input, low quality DNA. Here we have described LI-Seq, a method for cost-effective WGS library preparation that can be used for both high quality, high input and low quality, low input samples. Results suggest that, with the modifications described above, one can successfully produce high quality sequence data from DNA with input as low as 0.48 ng, for a fraction of the cost of traditional library preparation methods. More specifically with this method, we were able to prepare approximately 12 libraries for the same price as a single library with a more traditional WGS library preparation kit, thus allowing us to sequence more than an order of magnitude more samples. Overall, the increase in efficiency and cost-effectiveness provided by our method will allow conservation biologists to more broadly apply WGS methods to samples collected with non-invasive or minimally invasive methods.

Another potential challenge of using low input DNA with WGS methods is biased or incomplete amplification of the genome due to the low number of copies of the entire genome present (Meynert et al 2014). To assess this, we prepared a library from high quality, high input DNA (extracted from blood) in addition to preparing a library from low quality, low input DNA (extracted from feathers) using the same protocol. The final library molarity and fragment distribution was similar with both DNA sources, suggesting that the lower input DNA yielded equally high-quality libraries as high input DNA. In addition to impacting the quality of the final library, low input DNA could result in biased amplification leading to preferential sequencing of certain regions of the genome and result in less equal coverage across the genome than with high input DNA. The observed correspondence in patterns of genome-wide coverage between the libraries is to be expected in the absence of external influences causing biases related to the quality of input DNA (Ekblom, Smeds, and Ellegren 2014). Additionally, we checked for differential GC bias between the two libraries because it can indicate that biases were introduced during library preparation (Sims et al 2014). Our breadth of genome coverage (i.e. proportion of the genome sequenced) of 64% and 66% for the two libraries (high input and low input respectively) is comparable to that of mammalian genomes sequenced at similar depth (Green 2007). This breadth of coverage is suitable for numerous conservation genomic applications (e.g. identifying inbreeding across the genome) and resource efficient considering that approximately 30x coverage depth is required to achieve 95% coverage breadth (Sims et al 2014). Overall, our results suggest that when our method is employed, libraries prepared from low input and high input DNA both produce high quality sequence data.

When applying this protocol to other species and sample types, researchers may want to think about a couple important considerations. First, the genome size of the study organism should be considered. The protocol presented herein is an excellent option for use with organisms with a small genome, as the protocol was optimized using DNA from birds which on average have a genome size of 1.1 Gb. For species with larger genomes, more sequencing is required to achieve the same level of coverage, and therefore may require optimized methods and will have a relatively higher cost as well. Additionally, the ratio of tagmentation enzyme to input DNA may need to be adjusted in order to maintain a similar average fragment size. The tagmentation enzyme is one of the most expensive components of this method, so increasing the amount of enzyme per sample could also increase the per sample cost. Second, when considering combining libraries from low quality, low input and high quality, high input samples into a single sequencing run, researchers may want to take additional steps to ensure equal representation of the libraries. Specifically, performing a double size selection on the individual libraries prior to quantification and pooling can help ensure that the most accurate concentrations are used when pooling individuals for sequencing and, therefore, the most equal sequencing effort per sample will occur (Zamudio et al unpublished). Overall, with the aforementioned modifications taken into account, the method presented here could be applied more broadly to increase the efficiency and cost-effectiveness of WGS across a multitude of taxa.

Here we present a cost-effective method for producing WGS libraries using low input DNA from minimally-invasively collected samples. LI-seq provides a much-needed tool to bridge the gap between the conservation management applications of WGS data and frequently collected sample types, such as feathers and other non-invasively collected samples. Although a recent method was published that also provided a method for preparing non-invasively collected samples for WGS (Taylor et al 2020), the per sample cost may be prohibitive for use with population-scale studies for conservation efforts. By providing an efficient, cost-effective WGS method for low quantity and quality DNA samples, we hope conservation management efforts will be able to better take advantage of the applications WGS can provide for enhancing management efforts.

## Supporting information

Supplemental Full Protocol

## Acknowledgements

Thank you to Dr. Nina Therkildsen fo sharing her WGS method, Harmony B. Borchardt-Wier for early guidance on trouble-shooting Dr. Therkildsen’s protocol, and Dr. Cristian Gruppi for his technical advising. We thank the UC Davis Genome Center for their help with the sequencing. This work was made possible by an NSF CAREER award (008933-00002), an NSF grant Rules of Life grant (007604-00002), and a National Geographic grant (WW-202R-170) to K. Ruegg as well as the Extreme Science and Engineering Discovery Environment (XSEDE) computer cluster, which is supported by National Science Foundation grant ACI-1548562.

## Author Contributions

TMS and KCR designed the experiments. TMS carried out the experiments. MGD and KCR analyzed data. TMS, MGD, and KCR wrote the manuscript.

