## Supplemental Full Protocol for "LI-Seq: A Cost-Effective, Low Input DNA method for Whole Genome Library Preparation"

### LI-Seq Library Prep Protocol

*2021 Version 2: Added additional indices, changed protocol name to LI-Seq.*

***Method optimized by Ruegg Lab and the Bird Genoscape Project at Colorado State University; publication pending; please do not share without written permission from****

| **Reagents** | **Supplier** | **Cat. #** |
|---|---|---|
| Nextera XT index kit v2 Set A and/or D | Illumina | FC-131-2001/4 |
| Tagment DNA TDE1 Enzyme and Buffer Kit | Illumina | 20034198 |
| Kapa HiFi Kit | Kapa Biosystems | KK2612 |
| AMPure XP Beads* | Beckman Coulter | A63880 |
| Primer P1, 25nmol, purify NGS grade HPSF *(AATGATACGGCGACCACCGA)* | Eurofins | N/A |
| Primer P2, 25nmol, purify NGS grade HPSF *(CAAGCAGAAGACGGCATACGA)* | Eurofins | N/A |
| Ultrapure DNAse/RNAse-free Water | Thermo Fisher | 10977015 |
| Qubit dsDNA HS Assay Kit | Invitrogen | Q32854 |
| Qubit dsDNA BR Assay Kit | Invitrogen | Q32853 |

**Or Serapure beads*
***To sequence up to 384 samples together, purchase Set A and Set D for full complement*

**Important Notes Before Starting**

1. When working with such small volumes, evaporation is a major concern. Work quickly and keep tubes or plates sealed as much as possible.
2. Thorough mixing is very important at *every* step!
3. Make a working solution of your Primer P1 and Primer P2 at 10µM. (Primers are typically shipped dry and, with appropriate water added, become 100µM stock).
4. Use ultrapure DNAse/RNAse-free $H_20$ for all steps (normalization, 70% ethanol, etc).
5. After DNA normalization, steps 2 through 4 should be completed without pausing or freezing.
6. This protocol suggests a cleanup using Ampure beads on the individual libraries followed by a double size selection on the pooled library. It is possible to do double size selection on the individual libraries and save this second step. However, you may end up with wider fragment distribution depending on how precisely you size select. Alternatively, you can use a Pippin to size select the final library.
7. We recommend aliquoting indices, TDE1, and TD Buffer into strip tubes to decrease degradation by repeated freeze-thaw cycles since they are not intended to be used in such small volumes by Illumina. For indexes, it also it allows for quicker pipetting and decreases chances of accidentally swapping indices.
8. Reagents from steps 1 through 4 can be thawed on ice at the same time, usually takes about 45 minutes.
9. This protocol is written for using plates for library prep, but 8-strip PCR tubes can also be used.

#### 1 Normalize Genomic DNA (*estimated time: 2 hours*)

##### Preparation

1. For extractions from blood or tissue, check gDNA quality by running a 2% Agarose gel and quantity by using Qubit.
2. Select the highest quality samples for the library prep.
   *Highest quality is defined as clean, high molecular weight bands on a gel. Smearing indicates DNA degradation and samples likely won't result in high quality libraries. Use these samples only if absolutely needed.*
3. For feather extractions, elute into 400 µL and concentrate to 10uL with Ampure or Serapure beads to maximize yield. You can skip a gel check but still quantify using Qubit.

##### Procedure

1. Quantify all samples using 2 µL on Qubit or equivalent fluorescence plate reader. Do not freeze samples in between quantification and normalization as sample concentrations may change as a result of freezing.
2. Prepare at least 10 µL of gDNA at a concentration of 2.5 ng/µL in a plate. Use HPLC water to dilute samples. For feather extractions, a range as low as 0.5 ng/µL - 5.0 ng/µL has been tested with success. Feather samples below 0.5 ng/µL may require duplicate library preparation.
3. Re-quantify normalized samples with Qubit HS Kit. The concentrations won't be perfect but for blood or tissue extractions, a range from 1.0 ng/µL – 5.0 ng/µL is acceptable. For feather samples below 2.5 ng/µL, it is not necessary to quantify them again.
4. Place samples in 4$^{o}$C until use, unless it will be greater than a week. If normalized samples are frozen prior to tagmentation, quantify again to confirm that concentration is still in acceptable range.

#### 2 Tagmentation of Genomic DNA (*estimated time: 1 hour*)

##### Preparation

5. Remove the TD buffer and TDE1 enzyme from the -20ºC freezer and thaw on ice.
6. Remove normalized gDNA from 4ºC or freezer.
7. After fully thawed (20-40 min), mix TD buffer and gDNA by gently vortexing. Spin down.
8. Mix the TDE1 enzyme by pipetting up and down 20 times \. Do not vortex!

##### Procedure

9. Make the Tagmentation Master Mix (TMM) by mixing TD Buffer and TDE1 enzyme in a tube. Mix thoroughly by gently pipetting the mixture up and down 20 times. Use this reaction ratio to calculate the amount of each reagent needed.

| | 1x Tagmentation Master Mix (µL) |
|---|---|
| TD Buffer | 2.50 |
| TDE1 Enzyme | 0.50 |
| **Total Per Well** | **3.00** |

10. Include 10% pipetting error in your calculation.
    *For example, if you have 30 samples: 30 samples + 10% error = 33 samples. So calculate reagent volumes for 33 samples.*

11. Distribute TMM into 96-well plate. Either individually pipette 3.0 µL per well or a reservoir to pipette with a multi-channel.
    *It is highly recommended to pipette individually. Working with such small volumes, you have a higher risk of losing TMM by working a multi-channel and strip tubes.*
12. With a multichannel pipette, transfer 1 µL of well mixed gDNA into the tagmentation plate and gently pipette up and down 10 times. Change tips after every transfer.
    *Final reaction volume per well is 4.0 µL.*
13. Cover the plate with seal.
14. Give the plate a quick spin to collect all liquid at the bottom. Check that there are no large bubbles.
    *Bubbles can inhibit proper mixing of reagents and DNA and result in low yield. To remove bubbles, gently flick bottom of wells until bubble pops. Then centrifuge briefly again to collect all liquid at bottom.*
15. Place the plate in the thermocycler and run the following program:

| Step | Temperature | Time |
|---|---|---|
| Incubation | 55 ºC | 20 min |
| Hold | 10 ºC | ∞ |

16. Let plate cool to 10 ºC. Proceed immediately to the next step.

#### 3 Indexing PCR (*estimated time: 2 hours*)

##### Preparation

17. Remove the KAPA HiFi Hot Start Master Mix (KMM) and the indices from the –20 °C and thaw on ice.

    *For easier use with multichannels, store the indices as aliquots in strip tubes.*

18. After thawing, mix all reagents and indices by vortexing. Spin down briefly before opening.

##### Procedure

19. Each tagmented DNA sample (from Step 2) will receive an i7 Illumina index, an i5 Illumina index, and Kappa Master Mix. Each sample will then have a unique pair of dual indexes.
20. Typical kits can index up to 96 or 384 samples. Below is a map of how to visualize what sample would receive what indices in a 96-well plate.

| | | Set A i7 Indices | | | | | | | | | | | | Set D i7 Indices | | | | | | | | | | | |
|---|---|---|---|---|---|---|---|---|---|---|---|---|---|---|---|---|---|---|---|---|---|---|---|---|---|
| | | N701 | N702 | N703 | N704 | N705 | N706 | N707 | N708 | N709 | N710 | N711 | N712 | N716 | N718 | N719 | N720 | N721 | N722 | N723 | N724 | N726 | N727 | N728 | N729 |
| Set A i5 Indices | S502 | A1 | A2 | A3 | A4 | A5 | A6 | A7 | A8 | A9 | A10 | A11 | A12 | A1 | A2 | A3 | A4 | A5 | A6 | A7 | A8 | A9 | A10 | A11 | A12 |
| | S503 | B1 | B2 | B3 | B4 | B5 | B6 | B7 | B8 | B9 | B10 | B11 | B12 | B1 | B2 | B3 | B4 | B5 | B6 | B7 | B8 | B9 | B10 | B11 | B12 |
| | S504 | C1 | C2 | C3 | C4 | C5 | C6 | C7 | C8 | C9 | C10 | C11 | C12 | C1 | C2 | C3 | C4 | C5 | C6 | C7 | C8 | C9 | C10 | C11 | C12 |
| | S505 | D1 | D2 | D3 | D4 | D5 | D6 | D7 | D8 | D9 | D10 | D11 | D12 | D1 | D2 | D3 | D4 | D5 | D6 | D7 | D8 | D9 | D10 | D11 | D12 |
| | S506 | E1 | E2 | E3 | E4 | E5 | E6 | E7 | E8 | E9 | E10 | E11 | E12 | E1 | E2 | E3 | E4 | E5 | E6 | E7 | E8 | E9 | E10 | E11 | E12 |
| | S507 | F1 | F2 | F3 | F4 | F5 | F6 | F7 | F8 | F9 | F10 | F11 | F12 | F1 | F2 | F3 | F4 | F5 | F6 | F7 | F8 | F9 | F10 | F11 | F12 |
| | S508 | G1 | G2 | G3 | G4 | G5 | G6 | G7 | G8 | G9 | G10 | G11 | G12 | G1 | G2 | G3 | G4 | G5 | G6 | G7 | G8 | G9 | G10 | G11 | G12 |
| | S517 | H1 | H2 | H3 | H4 | H5 | H6 | H7 | H8 | H9 | H10 | H11 | H12 | H1 | H2 | H3 | H4 | H5 | H6 | H7 | H8 | H9 | H10 | H11 | H12 |
| Set D i5 Indices | S513 | A1 | A2 | A3 | A4 | A5 | A6 | A7 | A8 | A9 | A10 | A11 | A12 | A1 | A2 | A3 | A4 | A5 | A6 | A7 | A8 | A9 | A10 | A11 | A12 |
| | S515 | B1 | B2 | B3 | B4 | B5 | B6 | B7 | B8 | B9 | B10 | B11 | B12 | B1 | B2 | B3 | B4 | B5 | B6 | B7 | B8 | B9 | B10 | B11 | B12 |
| | S516 | C1 | C2 | C3 | C4 | C5 | C6 | C7 | C8 | C9 | C10 | C11 | C12 | C1 | C2 | C3 | C4 | C5 | C6 | C7 | C8 | C9 | C10 | C11 | C12 |
| | S517 | D1 | D2 | D3 | D4 | D5 | D6 | D7 | D8 | D9 | D10 | D11 | D12 | D1 | D2 | D3 | D4 | D5 | D6 | D7 | D8 | D9 | D10 | D11 | D12 |
| | S518 | E1 | E2 | E3 | E4 | E5 | E6 | E7 | E8 | E9 | E10 | E11 | E12 | E1 | E2 | E3 | E4 | E5 | E6 | E7 | E8 | E9 | E10 | E11 | E12 |
| | S520 | F1 | F2 | F3 | F4 | F5 | F6 | F7 | F8 | F9 | F10 | F11 | F12 | F1 | F2 | F3 | F4 | F5 | F6 | F7 | F8 | F9 | F10 | F11 | F12 |
| | S521 | G1 | G2 | G3 | G4 | G5 | G6 | G7 | G8 | G9 | G10 | G11 | G12 | G1 | G2 | G3 | G4 | G5 | G6 | G7 | G8 | G9 | G10 | G11 | G12 |
| | S522 | H1 | H2 | H3 | H4 | H5 | H6 | H7 | H8 | H9 | H10 | H11 | H12 | H1 | H2 | H3 | H4 | H5 | H6 | H7 | H8 | H9 | H10 | H11 | H12 |

21. You can see that i7 (N7xxx) indexes are distributed down “columns” of samples and i5 (N5xxx) indexes are distributed across “rows.” The volume per well is as below:

| | **1x Index Mix (µL)** |
|---|---|
| Kapa Master Mix | 6.0 |
| i5 Primer | 1.0 |
| i7 Primer | 1.0 |
| **Total Per Well** | **8.0** |

22. The following directions assume that your indexes are in strip tubes.
23. With a multichannel pipette, add 1µL of correct i5 primer to the plate, so that each *row* receives the same N5xxx index. Change tips after every transfer.
24. With a multichannel pipette, add 1µL of correct i7 primer to the plate, so that each *column* has the same N7xxx index. Change tips after every transfer.
25. With a multichannel pipette, add 6µL of Kapa Master Mix and mix thoroughly by gently pipetting the mixture up and down 20 times. Change tips after every transfer. The final volume per well is 12.0 µL.

26. Cover plate well with seal.
27. Give the plate a quick spin to collect all liquid at the bottom.
28. Place the plate in the thermocycler and run the following program. The program takes ~25 minutes to run.

| # of Cycles | Step | Temperature | Time |
|---|---|---|---|
| 1x | | 72° | 3 min |
| 1x | | 98° | 2:45 min |
| 8x | *Denaturation* | 98° | 15 sec |
| | *Annealing* | 62° | 30 sec |
| | *Extension* | 72° | 30 sec |
| 1x | | 72° | 1 min |
| | | 4° | ∞ |

29. Proceed immediately to the next step.

#### 4 Booster PCR (*estimated time: 1 hour*)

##### Preparation

30. Thaw KAPA HiFi Hot Start Master Mix (KMM) and 10 µM working stocks of Primers P1 and P2.

##### Procedure

31. Each indexed sample from Step 3 receives the same Booster Master Mix. Vortex and spin down the KMM and Primers P1 and P2. Calculate the volume of Master Mix by adding 10% pipetting error to the number of indexed samples. Use the recipe below:

| | **1x Booster Master Mix (µL)** |
|---|---|
| Kapa Master Mix | 7.6 |
| Primer P1 | 1.6 |
| Primer P2 | 1.6 |
| HPLC Water | 4.4 |
| **Total Per Well** | **15.2** |

32. Vortex the Master Mix and spin down. Pipette 15.2 µL into each well of the plate (so the final PCR volume is 27.2 µL). Mix by gently pipetting up and down 10 times. Change tips after every transfer.
33. Cover plate with seal.
34. Give the plate a quick spin to collect all liquid at the bottom (1000 rpm for 1 min)
35. Place the plate in the thermocycler and run the following program (Reconditioning PCR):

| **# of Cycles** | **Step** | **Temperature** | **Time** |
|---|---|---|---|
| 1x | | 95° | 5 min |
| 4x | *Denaturation* | 98° | 20 sec |
| | *Annealing* | 62° | 20 sec |
| | *Extension* | 72° | 2 min |
| 1x | | 72° | 2 min |
| | | 4° | ∞ |

36. You can safely pause and freeze or refrigerate the libraries at this point.

#### 5 Ampure Bead Size Selection (*estimated time: 2 hours*)

##### Preparation

37. Bring AMPure XP beads to room temperature (approximately 30 min). Protect from light while coming to room temp.
38. Prepare fresh 80% ethanol from molecular grade 200 proof ethanol in a sterile reservoir. You will need the # samples X 400 µL of 80% ethanol for washing. Use HPLC H2O to prepare and prepare 80% ethanol daily.

##### Procedure

39. Centrifuge the plate to collect all liquid (1000 rpm for 1 min.)
40. Vortex beads for 30 sec to ensure that they are evenly dispersed.
41. Perform “Left side selection” to remove small fragments (this is what is also referred to as a “bead cleanup”) using a ratio of 0.7:1. To DNA wells with 27.2 µL volume, add 19.0 µL of beads. Mix well by gently pipetting up and down 20 times. The color of the mixture should appear homogenous after mixing. Change tips in between samples.

    *Thorough mixing at this step is critical! Beads must physically interact with DNA in order to bind so thorough mixing increases the chances of those interactions. Beads are extremely high fidelity (1 µL of beads can bind up to 7 µg of DNA) so it is primarily the rate of interaction between beads and DNA that will limit DNA yield.*

    **Figure 1** Agilent High Sensitivity DNA chip Electropherogram.

    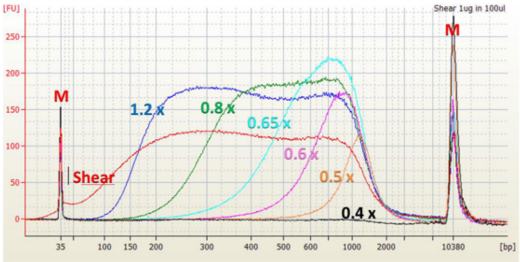
    

    *The figure above shows a “Left side selection” which is getting rid of small fragments at the left of the electropherogram. You can see that the larger the ratio of beads to DNA, the more small fragments are retained. Figure from Reference 4.*
42. Incubate at room temperature for 5 min. Desired DNA fragments are now attached to the beads.
43. Place the plate on the magnetic stand and incubate for about 2 min to separate beads from solution.

    *If solution is not clear after 2 min, give it more time. Wait until solution is completely clear before proceeding to next step.*
44. While the plate is on the magnetic stand, aspirate clear solution from the plate and discard. Do not disturb the beads. If the beads are accidentally pipetted, resuspend them back, wait for the solution to clear up, and repeat.

    *Small DNA fragments below ~320bp remain in solution and are discarded at this time.*

45. With plate still on the magnetic stand, dispense 200 µL of 80% ethanol into each well and incubate for 30 seconds at room temperature. Aspirate out ethanol without disturbing the beads and discard. Repeat for a total of 2 washes.
    *If for any reason your samples may have higher salt concentration, you can add a 3rd ethanol wash. This will slightly decrease DNA yield but increase purity.*
46. Remove the remaining ethanol with a 10 µL pipette.
47. Let the plate air dry for approximately 1-2 min (maximum of 5 min). Do not over dry the beads. This will result in lower yield as DNA will remain bound to over dry beads and not resuspend into solution.
    *Beads should appear glossy but not overly wet. No large ethanol drops should remain. If they do, use a pipette to remove them as opposed to waiting to evaporate. If beads turn rusty colored or appear dry or cracked, immediately proceed to the next step.*
48. Take the plate off the magnetic stand. Add 32 µL of 1mM Tris-HCl (pH 8) to each well of the plate. Carefully resuspend the beads by mixing 10-15 times. Incubate for 5 min at room temperature. DNA is now in the solution.
49. Place the plate back onto the magnetic stand and incubate for about 2 min to separate beads from solution.
    *If solution is not clear after 2 min, give it more time. Wait until solution is completely clear before proceeding to next step.*
50. While the plate is on the magnetic stand, aspirate 30 µL from the plate and transfer to a fresh plate. Do not disturb the beads. If the beads are accidentally pipetted, resuspend them back, wait for the solution to clear up, and repeat. Change tips after each transfer.
51. You can safely pause and freeze or refrigerate the libraries at this point.

#### 6 Pooling Libraries

##### Preparation

52. Qubit all samples. If any final library is below 1 ng/µL, reprep starting at the tagmentation step and use the DNA from the starting DNA dilution plate.
53. In order to have equal sequencing effort across individual libraries, samples must be pooled to equal number of copies. In this case, since all libraries should have the same average library size, simply use concentration to determine how much to add.

##### Procedure

54. Determine the library with the lowest concentration and see what the maximum amount of DNA (concentration in ng/µL x volume) that can be added from that sample. Use that value to calculate the volume of other samples to add so that each has the same amount of DNA present in final pool.

    *For example: Sample 1 is 10.3 ng/µL and you have 20 uL of it. Sample 2 is 3.7 ng/µL and you have 20 µL of it. The amount of DNA in Sample 2 is 74 ng (=3.7 ng/µL x 20 µL). In order to have the same amount of DNA from sample 1, you also need to add 74 ng of sample 1. Therefore, you would add 7.18 µL (=74 ng / 10.3 ng/µL). Therefore, the pool of these two samples now has 74 ng of each.*
55. Vortex samples to mix.
56. Centrifuge the plate to collect all liquid (1000 rpm for 1 min.)
57. Place the plate on magnetic stand, add volumes of samples as calculated.

#### 7 Ampure Bead Double Size Selection on Final Library

##### Preparation

58. Bring AMPure XP beads to room temperature (approximately 30 min). Protect from light while coming to room temp.
59. Prepare fresh 80% ethanol from molecular grade 200 proof ethanol in a sterile conical tube. You will need the # samples X 2000 µL of 80% ethanol for washing. Use HPLC H2O to prepare and prepare 80% ethanol daily.
60. Note that a double size selection using Ampure beads requires precision. The two most important parts are to a) pipette bead volumes accurately by using appropriate pipette and wiping droplets of beads from pipette tip exterior, and b) not transferring beads with solution in any step. We are selecting for a pretty narrow range (320-500bp) and adding too little or too much bead can shift that range dramatically.

##### Procedure

61. Vortex and centrifuge the pooled library to collect all liquid.
62. Vortex beads for 30 sec to ensure that they are evenly dispersed.
63. First, perform “Right Side Selection” to remove large fragments using a ratio of 0.63:1 (which will remove above ~600bp). Calculate the volume of beads to add by multiplying the volume of your pooled library times 0.63. Add that volume of AMPure beads into 1.5 mL tube with pooled library.

    *Do not just rely on your calculated pooled library volume. Use a pipette to accurately measure the volume! Also, make sure that you can add the full volume of beads from the left and right size selection (=0.73 x volume of library) to your tube size!*

64. Mix well by gently pipetting up and down 20 times or vortex for 15 sec. The color of the mixture should appear homogenous after mixing. Change tips in between samples.
65. Incubate at room temperature for 10 min. Large DNA fragments are now attached to the beads.
66. Place the plate on the magnetic stand and incubate for 5 min to separate beads from solution.
    *If solution is not clear after 5 min, give it more time. Wait until solution is completely clear before proceeding to next step.*
67. While the plate is on the magnetic stand, aspirate clear solution from the tube and transfer to new tube. **<u>Do NOT discard supernatant!</u>** This has the desired DNA. Transfer the entire volume minus 10 µL of supernatant without disturbing the beads. If the beads are accidentally pipetted, resuspend them back, wait for the solution to clear up, and repeat.
    *It is very important to avoid transferring beads at this step. If you do, you may end up with a peak in large fragment range during fragment analysis. If this is your first time performing a double size selection, consider sealing the tubes with first beads and save until after you've confirmed appropriate fragment distribution of samples.*
68. Now, perform "Left side selection" to remove small fragments (this is what is also referred to as a "bead cleanup") using a ratio of 0.73:1. To calculate how much to add, multiply your original library volume x 0.1 (which is the difference between the ratios of the two selections, 0.73-0.63).
69. To supernatant in new tube, add that many µL of beads. Mix well by gently pipetting up and down 20 times. The color of the mixture should appear homogenous after mixing. Change tips in between samples.
    *Note that the volume of beads is not equal to 0.73 x library µL. To calculate the volume of beads to add in the left side selection, first subtract the bead ratios (in this case, 0.73 – 0.63 = 0.1). Use the calculated value (0.1) x the starting volume of sample to calculate the volume of beads to add. This is because it is the ratio of salts in solution more than the beads itself that dictate what fragment sizes are bound to beads. Since the supernatant from the first beads contains all of the reagents from those beads, we subtract the ratios to get the final value. However, because we are adding a relatively small volume of beads to a larger supernatant volume, <u>thorough mixing at this step is critical</u>! Beads are extremely high fidelity (1 µL of beads can bind up to 7 µg of DNA) so it is primarily the rate of interaction between beads and DNA that will limit yield.*
70. Incubate at room temperature for 10 min. Desired DNA fragments are now attached to the beads.
71. Place the plate on the magnetic stand and incubate for about 5 min to separate beads from solution.
    *If solution is not clear after 5 min, give it more time. Wait until solution is completely clear before proceeding to next step.*

72. While the plate is on the magnetic stand, aspirate clear solution from the tube and discard. Do not disturb the beads. If the beads are accidentally pipetted, resuspend them back, wait for the solution to clear up, and repeat.
    *Small DNA fragments below ~320bp remain in solution and are discarded at this time.*
73. With plate still on the magnetic stand, dispense 1000 µL of 80% ethanol into each well and incubate for 30 seconds at room temperature. Aspirate out ethanol without disturbing the beads and discard. Repeat for a total of 2 washes.
74. Remove the remaining ethanol with a 10 µL pipette.
75. Let the tube air dry for up to 5 min. Do not over dry the beads. This will result in lower yield as DNA will remain bound to over dry beads and not resuspend into solution.
76. Take the tube off the magnetic stand. Add xx µL of 10mM Tris-HCl (pH 8). Carefully resuspend the beads by mixing 10-15 times. Incubate for 10 min at room temperature. DNA is now in the solution.
    The volume you elute into will depend on two factors: 1. Your sequencing platform and 2. The number of lanes you are running. Confirm with your sequencing core on sample requirements prior to completing this step. You can always add additional volume via dilution.
77. Place the tube back onto the magnetic stand and incubate for about 5 min to separate beads from solution.
    *If solution is not clear after 5 min, give it more time. Wait until solution is completely clear before proceeding to next step.*
78. While the tube is on the magnetic stand, aspirate 2 µL less then the volume you added to the tube and transfer to a fresh tube. Do not disturb the beads. If the beads are accidentally pipetted, resuspend them back, wait for the solution to clear up, and repeat.

#### 8 Final Library QC

##### Procedure

79. Qubit the final concentrated pool and run on fragment analyzer (such as bioanalyzer or tapestation). Typically, high sensitivity kits work best (but you'll need to dilute the library concentration to ~2 ng/µL).
80. If fragment analysis looks good (with fragment range ~320-600bp), library is ready for submission. Confirm with your sequencing core what volume, concentration, and insert size requirements are for your library!

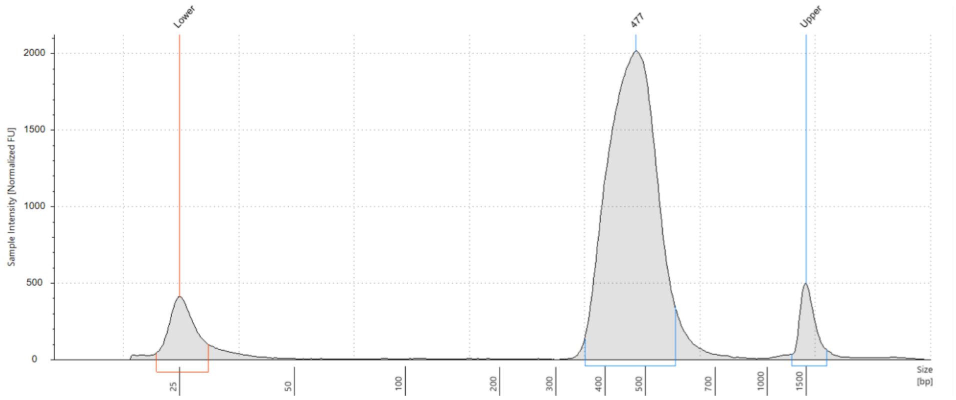


*Example of a final library QC after Pippin size selection. Note that size selection is never perfect but removing fragments below 320bp is most important for quality sequencing.*

#### 9 Optional Reconditioning PCR

If fragment analysis shows signs of overamplification or final library is too low molarity, perform an additional reconditioning PCR at 1 cycle. Use 12 µL of final pooled library and follow same steps as Part 4, but only run for 1 cycle in thermocycler. Follow with bead cleanup from Step 7.
